# Lipid-Uptake Pathways and Lipid-Protein interactions in P-glycoprotein Revealed by Coarse-Grained Molecular Dynamics Simulations

**DOI:** 10.1101/191239

**Authors:** E. Barreto-Ojeda, V. Corradi, R.-X. Gu, D.P. Tieleman

**Affiliations:** Department of Biological Sciences and Centre for Molecular Simulation University of Calgary, 2500 University Dr. NW, Calgary, AB T2N1N4, Canada

## Abstract

P-glycoprotein (P-gp) exports a broad range of dissimilar compounds, including drugs, lipids and lipid-like molecules. Due to its substrate promiscuity, P-gp is a key player in the development of cancer multidrug resistance (MDR). Although P-gp is one of the most studied members of ABC-transporters, the mechanism of how its substrates access the cavity remains unclear. In this work, we performed coarse-grained (CG) molecular dynamics (MD) simulations to explore possible pathways of lipid-uptake in the inward-facing conformation of P-gp embedded in bilayers with different PC:PE lipid ratios. Our results show that in the inward facing orientation only lipids from the lower leaflet are taken up by the transporter. We identify positively charged residues at the portals of P-gp that favor lipid entrance to the cavity, as well as lipid binding sites, in good agreement with previous experimental studies. Our results show no selectivity for PC vs. PE lipids. We offer several examples of lipid uptake-pathways for PC and PE lipids that help to elucidate the molecular mechanism of substrate-uptake in P-gp.

## Introduction

P-glycoprotein (P-gp), a member of the ATP-Binding Cassette (ABC) transporters superfamily, is responsible for the translocation of a wide range of compounds across the plasma membrane [1][2]. P-gp is found in several tissues facing an excretory compartment, for example, in the apical surface of epithelial cells of liver, pancreas, and small intestine [3]. It is also found in the brush border of kidneys and the endothelial cells of the blood brain barrier [4][5]. Natural substrates of P-gp are toxins and xenobiotics, against which P-gp provides a mechanism of defense by exporting them outside the cell. However, substrate binding to P-gp is highly promiscuous [6]–[8]. Its substrates, many of them structurally quite dissimilar compounds, include metabolites, drug molecules like analgesics and antibiotics, drug-like molecules such as diagnostic dyes, and lipids [9]–[13]. In addition, several studies have linked the over-expression of ABC transporters, and most notably P-gp, with mechanisms of resistance in various cancer cells [14]. Due to its broad substrate specificity, P-gp confers multidrug resistance (MDR) by pumping several pharmacological agents to the extracellular medium, preventing their accumulation inside the cell [15][5]. The development of MDR is considered one of the major challenges in cancer treatment [11][16][17], and since P-gp acts as a key player, the study of its underlying mechanism is of considerable importance to improve current chemotherapeutic treatments [5][18].

In addition to the drug transporter function, P-gp has also been proposed to act as lipid flippase [19], flipping lipids from the inner to the outer leaflet [5]. Direct interaction between lipid-based molecules and P-gp have been identified in previous studies [12], [13]. Competition of transport and inhibition of ATPase activity suggest that lipids are not only active modulators of the transport mechanism of P-gp, but also substrates [20]. The high affinity of the transporter for lipids and lipid-based molecules as miltefosine and edelfosine, phospholipid-based anticancer agents, has been linked with the flippase model [5]. Given the crucial role in cancer MDR, further understanding of the factors that determine lipid binding is essential for targeting P-gp inhibition as a means to increased drug-delivery in chemotherapy treatments.

The P-gp structure consists of two units linked by a flexible linker. Each unit includes an N-terminal transmembrane domain (TMD), followed by a C-terminal nucleotide-binding domain (NBD). The TMDs, with six transmembrane helices each (TMD1, H1-H6; TMD2, H7-H12), form the translocation pathway for substrates. TMDs and NBDs are coupled by intracellular loops (ICLs). As for any other ABC transporter, the substrate translocation mechanism of P-gp is ATP driven [21]. ATP binding and hydrolysis at the NBDs induce NBD dimerization and dissociation, which in turn drive conformational changes in the TMDs [1][6]. To allow transport, P-gp switches between inward-facing and outward-facing states, relevant for substrate binding and release, respectively [5][16]. In the inward-facing conformation [22], the two TMDs (TMD1, TMD2) form a cavity open towards the cytoplasm, enclosed by helices H1, H3, H5, H7, H11 and H12. In addition, two portals allow access of molecules directly from the bilayer, Portal 1 (P1) and Portal 2 (P2). P1 is constituted by helices H4 and H6, and P2 by H10 and H12. Although several theoretical and experimental studies have targeted substrate binding [8–10], the mechanism by which P-gp can recognize substrates remains elusive.

As a step towards a better understanding of how substrates access the translocation cavity, in this study we use molecular dynamics (MD) simulations at a coarse-grained (CG) level of detail to investigate lipid uptake and distribution in P-gp using the Martini force field [25]. CG models are a powerful tool to investigate biomolecular processes of larger systems, or longer time scales, due to the simplified level of detail while maintaining chemical specificity [25]. CG models are particularly suitable to reach system size and time scales relevant to comprehensively study substrate pathways [26]. The Martini force field has been widely used to resolved lipid-protein interactions in diverse biomolecular systems [27] [28]. For example, Karo *et* al. explored the binding of mitochondrial creatine kinase (MtCK) to the membrane [29]. Sengupta *et* al. analyzed the cholesterol binding sites in the serotonin_1A_ receptor [30]. Schmidt *et* al. investigated phosphatidylinositolphosphate (PIP_2_) binding events in the potassium channel Kir2.2 [31].

Given the importance of the role of lipids as possible substrates in P-gp, we investigated lipid-protein interactions using the crystallographic structure of mouse P-gp in the inward-facing conformation [22] embedded in membranes consisting of different ratios of POPC and POPE lipids. To investigate the P-gp lipid-uptake we first analyzed lipid access to the cavity to evaluate a possible selectivity between PC and PE lipids. Secondly, we studied the interactions between lipids and protein residues at the portals and inside the cavity. Finally, we analyzed the dynamics of the lipids inside the cavity of P-gp and identified key interactions between the lipid head groups and the lipid tails with the residues in the cavity. In agreement with previous experimental reports, our results highlight the importance of positively charged residues at the portals, such as Arg355, that may facilitate the lipid-uptake via electrostatic interactions. We identify key binding residues in the cavity of P-gp, for lipid head groups and lipid tails, and display the lipid occupancy in the inner region of the cavity.

## Methods

We used the refined structure of mouse P-gp as input structure for MD simulations (PDB ID 4M1M) [22]. We conducted coarse-grained (CG) molecular dynamics (MD) simulations of P-gp embedded in lipid bilayers of different composition (Table 1) including pure POPC and POPE bilayers, a 1:1 symmetric POPC:POPE bilayer, and two asymmetric bilayers: one with the same PC:PE ratio as reported for the mammalian plasma membrane [32] and for comparison one with the PC:PE ratio inverted between the two leaflets. All simulations were performed using the GROMACS 4.6 package [33] and the standard Martini force field 2.2 for protein and 2.0 for lipids. [25][27][33][34]. Simulation time was 20 μs for each system.

**Table 1.**
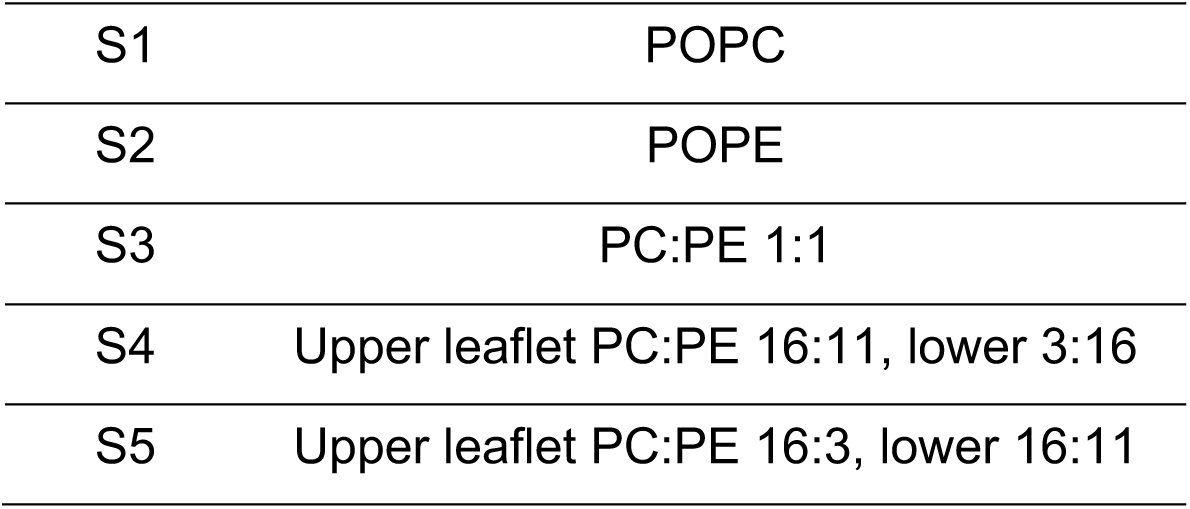
Lipid composition of the bilayers.

## Simulation Setup

The atomistic structure of P-gp was converted to the corresponding CG model using the Martinize [35] building tool. An elastic network was applied for atom pairs within a 1.0 nm cut off distance [36]. All protein-bilayer systems were built using the INSANE (INSert membrANE) CG tool [37]. Membranes were solvated with standard Martini water. Ions were also added to neutralize the protein and to reach a final concentration of 150 mM of NaCl. Each simulation box of 140 x 140 x 180 Å contains 1 copy of P-gp in CG representation and ~600 molecules of lipids, ~23000 CG water, ~250 Na^+^ ~260 Cl^-^.

Energy minimization for all systems was carried out using the steepest-descent algorithm with a force tolerance of 100 kJ mol^-1^ nm^-1^ for convergence. For electrostatic interactions, we used a shift function to turn off the interactions between 0 and 1.2 nm. The relative dielectric constant was set to *ε*_r_=15, as standard in non-polarizable Martini. All subsequent equilibration simulations were performed for 8 ns in the NPT ensemble with a 10 fs time step. The Berendsen barostat [38] kept the systems at 1 bar with a relaxation time constant T_p_=1.0 ps with semi-isotropic pressure coupling. The reference temperature was set at 310K and controlled by velocity rescaling thermostat [39] with a relaxation time constant T_T_=0.5 ps. Position restraints on the protein with a force constant of 1000 kJ mol^-1^nm^-2^ were applied gradually, first to the backbone beads and then together with the elastic network. Production runs were performed with a 20 fs integration time step. The Berendsen barostat [38] was used to maintain the systems at 1 bar with a relaxation time constant T_p_=5 ps. The reference temperature of 310 K was controlled by the velocity rescaling thermostat [39] with a relaxation time constant τ_T_=2 ps. The electrostatic interactions were treated as described for energy minimization.

## Lipid uptake and lipid-protein interaction analysis

### Lipid uptake

The main cavity is defined here as formed by the residues on the helices lining the inner portion of the P-gp cavity, i.e. side chain residues located along the helices H1, H3, H5 and H7, H11, H12. Residues on H2, H8-H9 (external helices) and H4/6 and H11/12 (portals) were excluded. A lipid was considered as accessing the cavity if the corresponding PO_4_ bead was found for more than 200ns within 8 Å of any of the side chain residues of the main cavity. Among different values, the 8 Å cut off was the optimum distance to evaluate uptake events with no overestimating. The window of 200ns guarantee the uptake events in consideration are not caused by thermal fluctuations. Numerical analyses were carried out using open-source Python Numpy and MDtraj libraries [39][40].

### Lipid occupancy inside the P-gp cavity

A contact between a lipid molecule and the cavity was counted if the corresponding PO_4_ bead was located within 5 Å from any of the residues in the cavity of P-gp. We will refer to this as the contact condition. The 5 Å cut off was chosen considering that the minimum distance found between lipids and residues, was 4 Å. For each system, the number of lipids that satisfy the contact condition at the same time was calculated, representing the number of lipids that remain inside the cavity of P-gp simultaneously. The restriction that a contact must be more than 200ns does not apply in order to know the number of lipid molecules able to occupy the cavity simultaneously. We also calculated the time during which a given number of lipid molecules occupy the cavity simultaneously.

## Lipid-protein interactions

### Portals

To elucidate the lipid-uptake pathways of P-gp, we performed lipid-protein interaction analysis to identify the residues at the portals of P-gp following the contact condition. The portals of the protein are delimited by helices H4/6 and H11/12. We count a binding event when the PO_4_ bead of a lipid molecule is located within 5 Å of the residues at the portals of P-gp. The residues here consider at the portals are: Lys230 to His241 of H4, Ser345 to Gly356 of H6, Leu875 to Lys883 of H10, and Tyr994 to Ser1002 of H12. The analysis of binding events was performed on the lipid molecules selected from the previously described lipid uptake. Lipid binding sites were identified by the contact time with lipid head groups over the last 15 μs of simulation time. Lipid binding sites were identified as the top 5 residues for which the contact condition is satisfied during the last 15 μs of simulation time. The maximum contact time registered was normalized to 1.0. A color map distinguishes three levels of interaction: a very high interaction corresponds to between 0.8 and 1 (red); high interactions are between 0.4 and 0.7 (magenta); below 0.4 the interactions are moderate (grey).

### Cavity of P-gp

Similar to the analysis of the lipid-protein interactions at the portals, a lipid molecule was considered bound if the corresponding PO_4_ bead was located within 5 Å of the residues at the cavity of P-gp for at least 200ns. The lipid binding sites and the levels of interaction follow the same definition as explained above for the top 10 residues, in addition to the those interacting exclusively with PC (blue) and PE (green).

### Lipid tails inside the cavity

To analyze the dynamics of the lipid head groups and tails inside the P-gp cavity, we considered the PO_4_ bead of each lipid and the last bead of each tail, i. e. C_4A_, C_4B_ beads. The lipid binding sites for each tail were identified by the contact condition. For each frame, we defined the middle plane of the membrane z’=0 as the average z coordinates between the upper and lower leaflet. Region I is defined as the highest region in the P-gp cavity, z’>0; Region II corresponds to the region between the lower leaflet and the middle plane of the bilayer, where z’<0. Measurements of the *z* coordinate for the phosphate (PO_4_) bead and the last bead of each tail (oleoyl tail, C_4A_ bead; palmitoyl tail, C_4B_ bead) were carried out.

## Occupancy Maps

Fractional lipid occupancy was calculated for POPE and POPC lipid types over the last 15 μs of each simulation system, using the VolMap plugin in VMD 1.9.1, with a grid resolution of 1 Å [42]. For the systems with pure popc or pope bilayers maximum occupancy was found at 0.78 and 0.72, respectively. The maximum occupancy was found at 0.73 for the system with POPC:POPE 1:1 ratio, and at 0.78 and 0.74 the asymmetric bilayer with the plasma ratio and the asymmetric system with the inverted plasma ratio, respectively. For visualization purposes, we display the isosurfaces at 20%, 50% and 70% of the maximum value, set at 0.72 for all the systems, and the maps are superimposed with the protein structure extracted from the last frame of each simulation.

## Results

Lipid uptake events were identified by the interaction between the lipid head group and residues of helices H1, H3, H5 and H7, H11, H12, following the contact condition (See Methods). The number of lipid-uptake events were determined by the number of lipid head groups accessing the main cavity, for both PC and PE lipid types. However, from our simulations we found that these events involve only lipid molecules from the lower leaflet. Although we initially hypothesized that P-gp showed selectivity for POPE vs. POPC, the results show no indication this is the case (Fig SI1), and in what follows we generally combine the results for both lipids from all the simulations systems.

The level of detail from the CG approach, together with the extensive sampling for lipid diffusion in our simulations, allow us to observe reversible lipid binding in the cavity of P-gp and lipid confinement inside the cavity of P-gp. Two examples of these findings are illustrated in Figure 1. On the left panel, at time 0μs (red), the lipid is outside the cavity; during the first microsecond of simulation it enters the cavity through P1, where it remains for less than 1 μs before leaving the cavity. After ca. 9 μs (white), the lipid enters again the cavity through P2, where it stays for the remaining simulation time up to 20 μs (blue). The right panel in Figure 1 shows a lipid molecule that remains for ca. 16 μs inside the cavity before diffusing outside.

**Fig 1.**
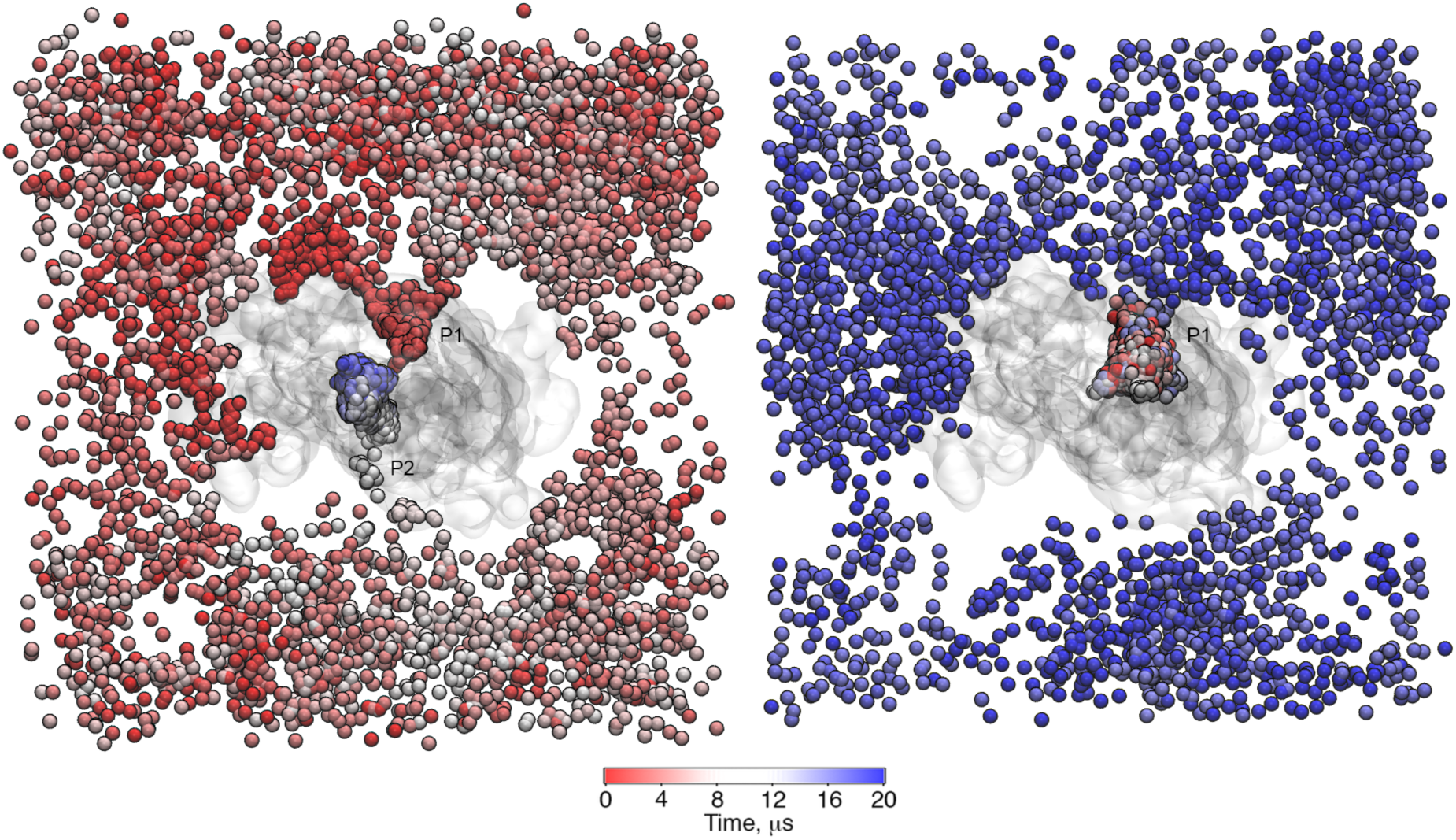
Two examples of PO_4_ bead trajectories found in the pure POPC bilayer. (Left panel) At time 0 μs, the lipid molecule is outside and permeates the cavity twice during the simulation time, during the first microseconds of the simulation through P1 and after ca. 9 μs through P2. **(Right panel)** The selected lipid molecule remains inside the cavity for ca. 16 μs before exiting through P2.

## Lipid occupancy inside the P-gp cavity

Using the contact definition described in the Methods, no PO_4_ beads of PC or PE lipids were found in the cavity for ca. 45% and ca. 60% of the simulation time, respectively (Figure 2A). Instead, water molecules were found inside. Overall, a single PC lipid remains in the cavity for ca. 40%, while a single PE remains during ca. 30% of the simulation time. Transiently, up to a maximum of four lipid molecules can occupy the cavity simultaneously, although for short periods of time (less than 0.05%), as shown in Figure 2A.

**Fig 2.**
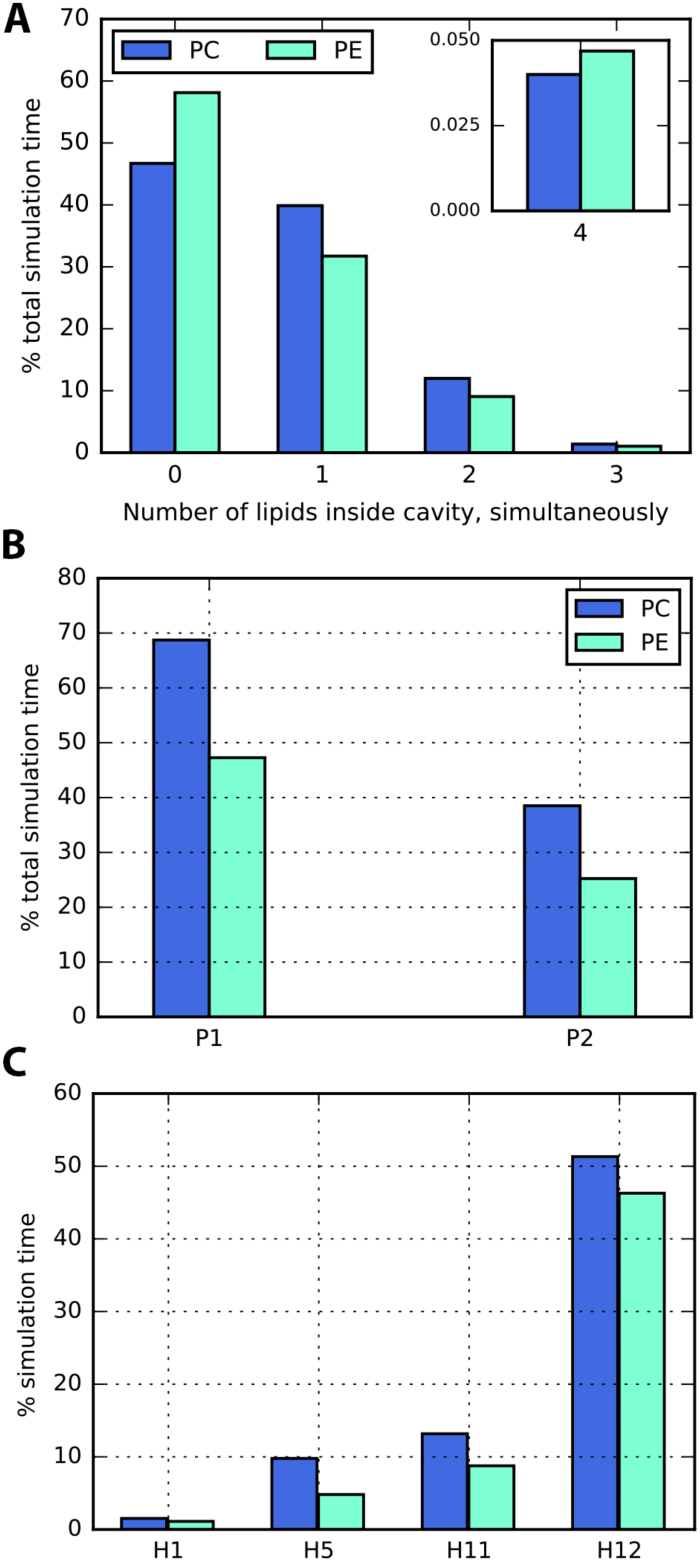
Lipid occupancy and lipid binding events in the cavity and at the portals of P-gp. Results are shown combining all the simulated systems. **(A)** The lipid occupancy in the cavity is primarily given by one lipid molecule for both PC and PE lipid types. Up to four lipid molecules can access the cavity simultaneously. **(B)** At P1, binding events for PC lipids were registered during ca. 70% of the total simulation time, PE binding events took place during ca. 45% of the time. Binding events at P2 are reduced to less than 40% for PC and ca. 35% for PE lipids. Overall, binding events occur for a longer period of time at P1 than at P2. **(C)** Lipid binding activity is detected at H12 during ca. 50% of the simulation time. Residues of H1/5/11 established lipid binding during less than 15% of the simulation time. In the cavity, binding events at H3/7/10 were not observed in our simulations.

## Lipid-protein interactions at the portals

To identify the binding sites for POPC or POPE lipids, we looked at the residues located at the portals (helices H4/6 at P1 and H11/12 at P2) that during the simulation time interacted with the head group of each lipid type. Every time a residue satisfied the contact condition, one binding event was counted (See Methods). As shown in Figure 2B, overall, P1 registered more binding events than P2 for both lipid types.

We then calculated the number of contacts between the PO_4_ beads to identify key binding residues at the portals, involved in the lipid-uptake. Despite the chemical difference between POPC and POPE head groups, the predominant binding sites found at the portals were the same for both lipid types: Arg355 located at H6, Tyr 994 at H12 (Figure 3). Additional interactions were registered with the residues Lys230, Lys235 at H4, Ser345 at H6, and Lys881, Asp993 at H12.

**Fig 3.**
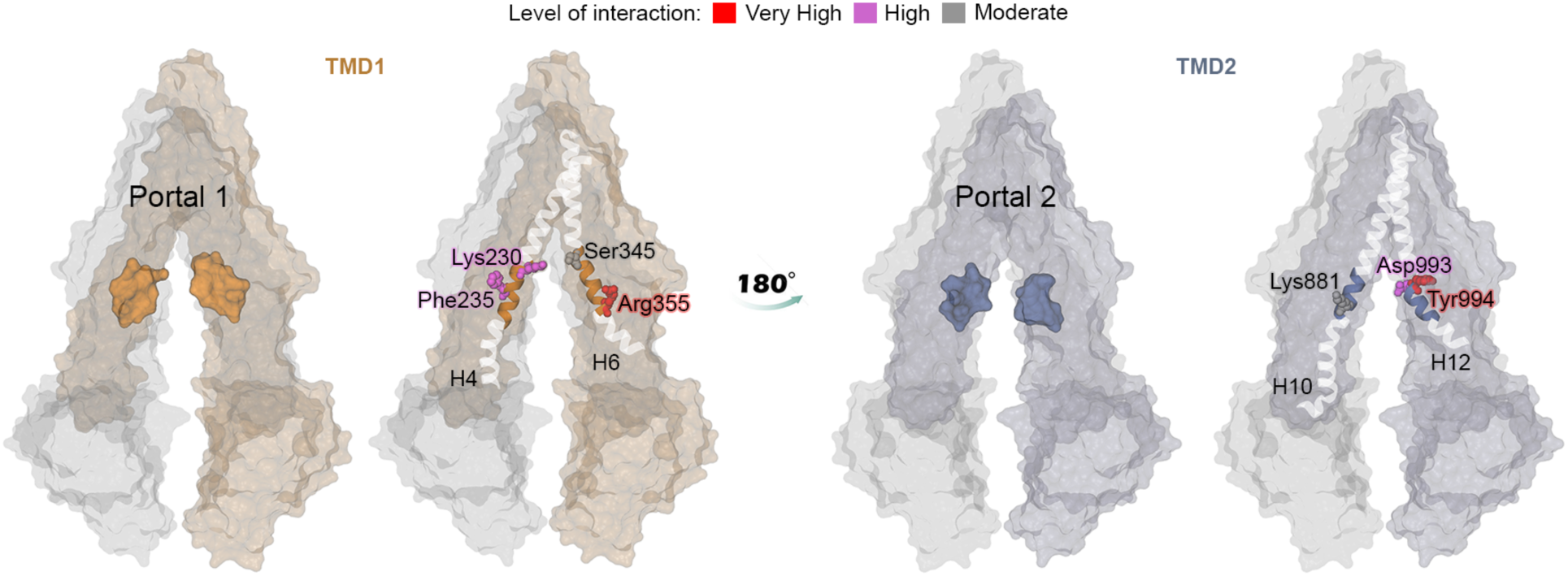
Binding residues for PC and PE lipids at the portals of P-gp. Binding residues are highlighted according to color map. The maximum contact time registered was normalized to 1.0. A color map distinguishes three levels of interaction: a very high interaction corresponds to between 0.8 and 1 (red); high interactions are between 0.4 and 0.7 (magenta); below 0.4 the interactions are moderate (grey). Lys 230, Phe235, Ser315, Arg355 at P1 and Asp993, Tyr 994 at P2 registered the highest 10% interactions.

## Lipid-protein interactions inside the cavity of P-gp

Similar to the previous analysis for the residues at the portals, we identified residues inside the cavity of P-gp involved in interactions with lipid molecules. PC and PE binding events mostly involved residues of H12, registered over ca. 50% of the simulation time (Figure 2C). PE lipid binding was observed over ca. 45% of the simulation time. Binding residues of H1/5/11 registered interactions during less than 15% of the time, while residues at H3, H7 and H10 did not show binding events over the course of the simulation. Inside the cavity, the lipid-protein interactions mainly involved the residues Val984, Val987 and Ser988 at H12, and residues Phe938 and Thr941 located along H11 (Figure 4B and SI2), all of them in the upper half region of the P-gp cavity. Additional binding events were found for residues His60, Ser294, Phe339 at the TMD1 (Figure 4A). We observed a very high similarity between the PC and PE binding residues. Only two binding residues showed selectivity for lipid binding: Ala338 for PC and Gly266 for PE, both with moderate interaction.

**Fig 4.**
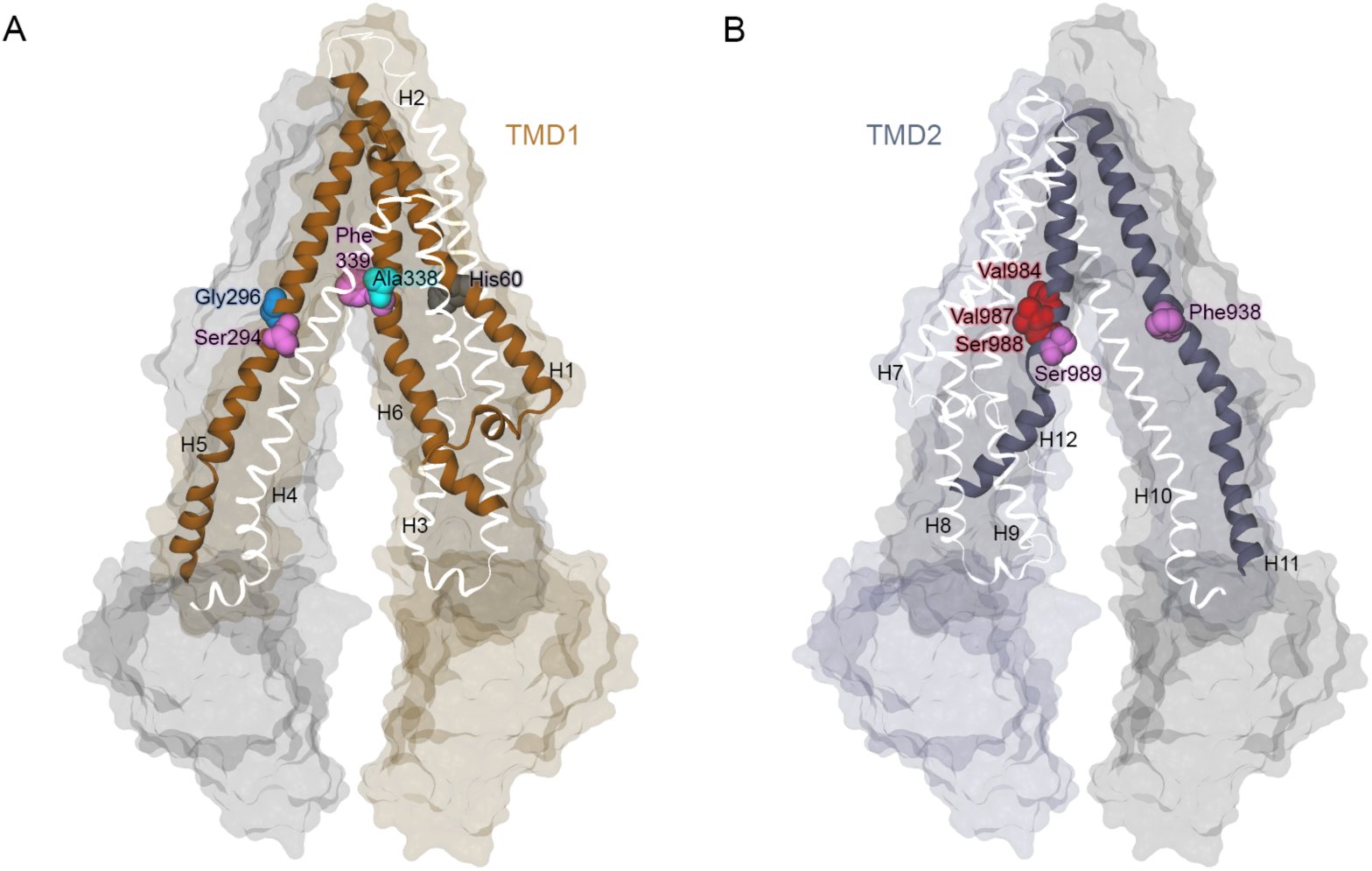
Binding residues in the cavity of P-gp. Residues Ser294, Phe339 of TMD1 **(A)** and Ser988, Val987, Val984 and Ser989 of TMD2 **(B)** are highlighted in the atomistic representation of P-gp. Gly296 and Ala338 registered interactions only with head groups of PE and PC, respectively. The color map follows the same scheme as in Fig 3.

## Dynamics of the lipid molecules in the P-gp cavity

To illustrate the dynamics of the PO_4_ bead and lipid tails inside the cavity of P-gp, we calculated the average z-coordinate over windows of 1 μs and plotted over the course of the simulation time (Figure 5B and 5D). In all our simulations, we observed that the PO_4_ bead remains mainly in the lower region of the membrane (Figure 5A and 5C) for both lipid types, PC and PE. The tails of the lipids, however, occupy distinct regions and extend to the upper region of the cavity, where oscillatory and flipping events were detected. We also identified the lipid tail binding sites in the upper most region of the bilayer, as established by the contact condition (see Methods). For the tails, we observed binding events mostly at H5 and to a lesser extent at H7. No binding residues were found on H1/3/11/12. Residues Phe299 and Leu301 showed very high interaction with the tails for both lipid types. A frequent interaction was identified between the oleoyl tail and Tyr303/Tyr306; and between the palmitoyl tail and Phe299/Ala304 (Figure SI3).

**Fig 5.**
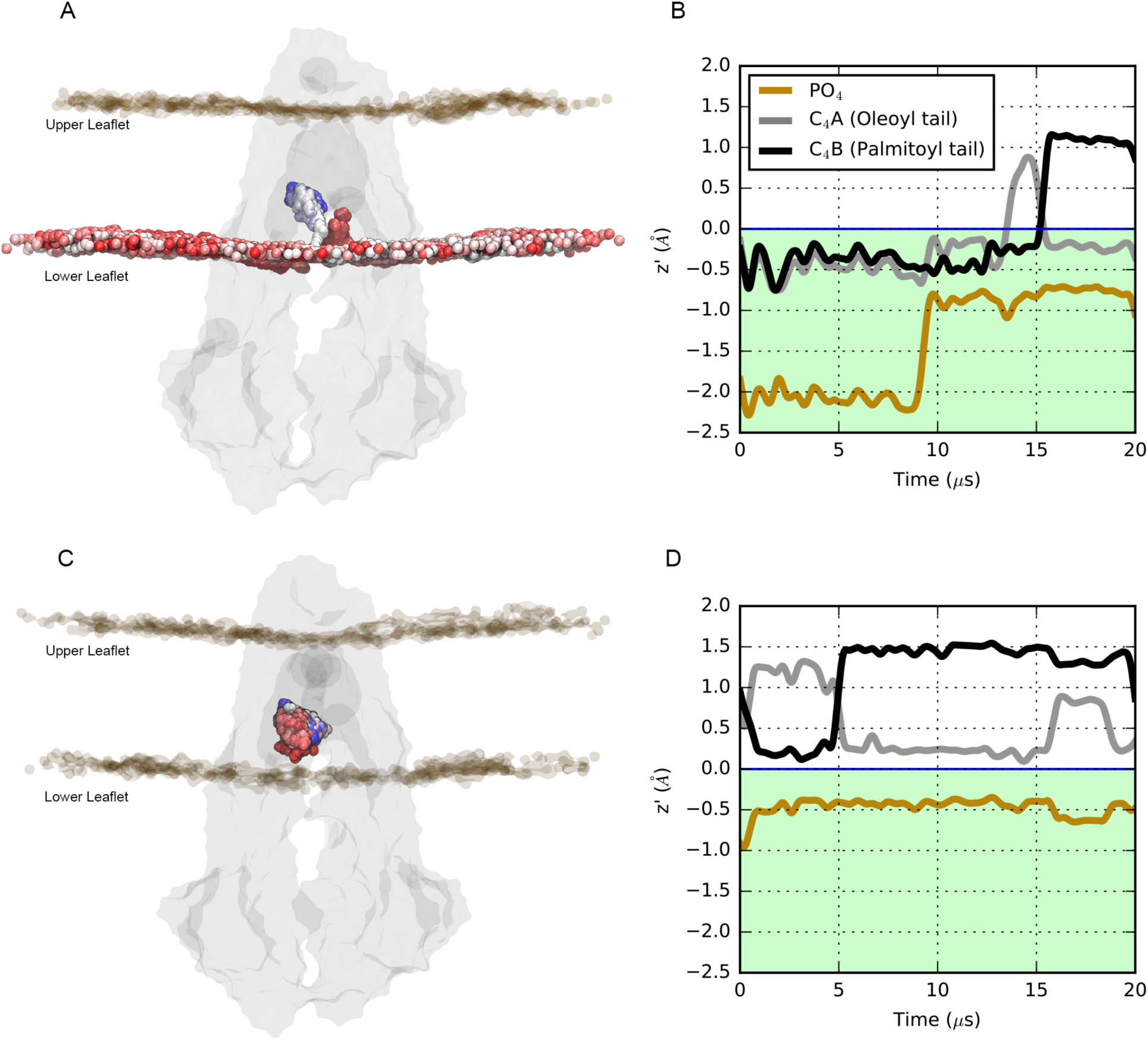
Dynamic of PO_4_ beads and tails inside the cavity of P-gp. The average z profile coordinate for the PO_4_ bead and tail beads (C_4A_ and C_4B_) plotted over 20 μs of simulation time for a lipid that experience two uptake events **(A-B)** and a lipid that remains in the cavity **(C-D)**. In **(A)** and **(C)**, P-gp is shown in gray surface, while the position of the selected PO_4_ bead is shown as colored sphere, from red to white to blue, as a function of time. The z profile of the PO_4_ bead (yellow line) and the C_4A_ and C_4B_ beads (grey and black lines) were calculated over windows of 1 μs.

The lipid occupancy maps highlight the ordering of the lipids in the immediate proximity of P-gp (see Methods). We observed high occupancy values inside the cavity, independent of the lipid composition (Figure 6). The presence of the lipids spans the entire available volume due to the lipid tails reaching the upper region, reinforcing the results showed in Figure 5. The lipid organization inside the cavity does not significantly differ between the PC:PE ratios considered in this work (Figures 6A and 6B). In addition, hot-spots of lipid occupancy are also detected at the portals, where lipids compete for access to the inner portion of the cavity.

**Figure 6.**
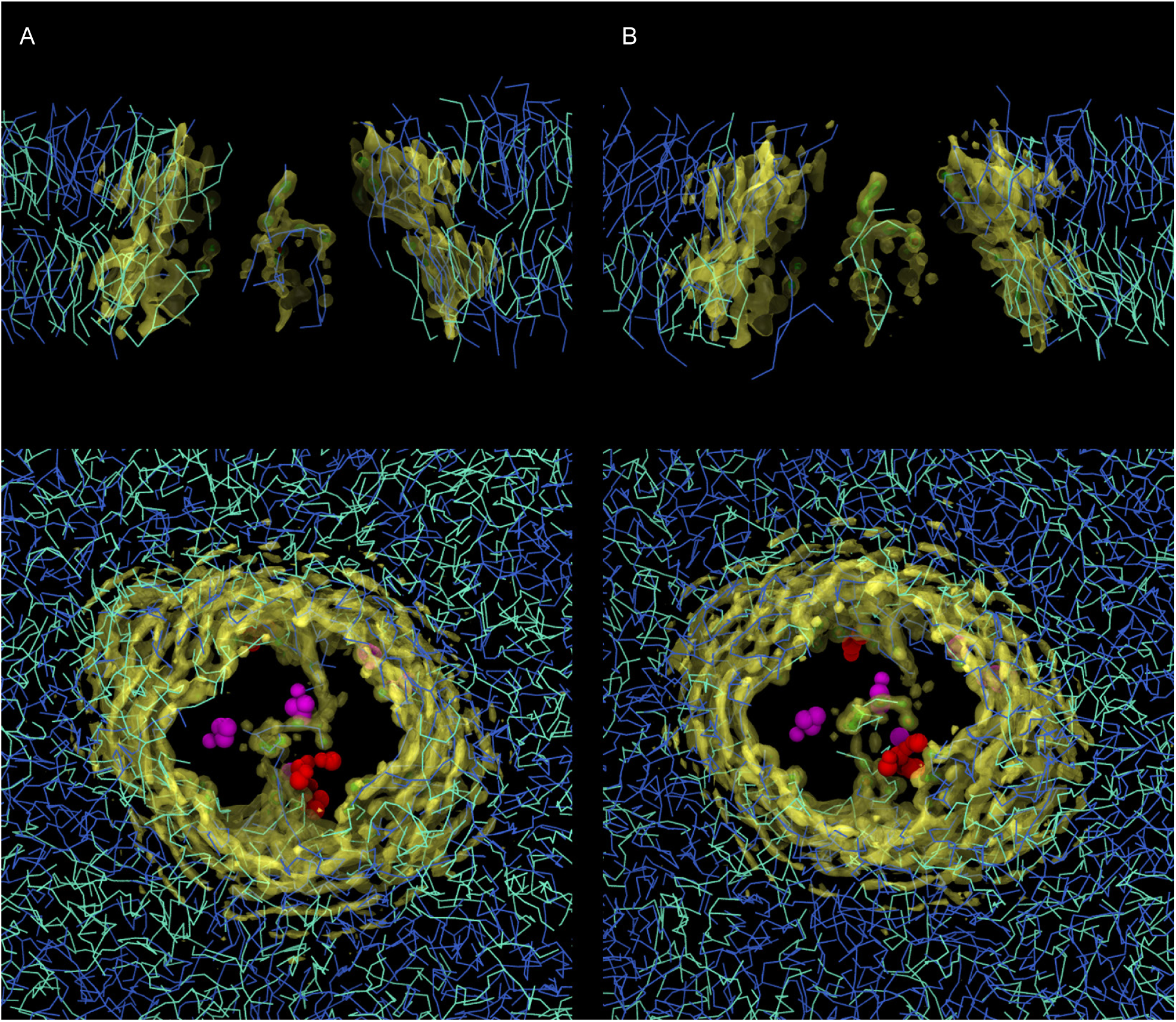
Lipid occupancy maps for the (A) POPC:POPE 1:1 and (B) plasma ratio membrane ratio PC:PE systems. The isosurfaces are shown for values corresponding to 20% (transparent yellow), 50% (transparent green), and 70% (red) of the maximum value. Top panel: Lateral view of the bilayer. The protein is omitted for clarity and the occupancy maps are clipped to highlight the occupancy signal in the inner region of the cavity. Bottom panel: Top view from the extracellular side. From the last frame of the corresponding simulation system, residues with a high degree of interaction are shown as magenta (Phe339 and Phe938) and red spheres (Val987, Ser988, Val984, Tyr994 and Arg355), following the same color scheme from Figure 3 and 4. PE and PC lipids are shown in cyan and blue lines, respectively.

## Lipid pathways

Our simulations enable us to identify trajectories of lipids that access the cavity of P-gp. Examples of trajectories of PC and PE lipid-uptake along 20μs of simulation time are illustrated in Figure 7. Simultaneous lipid uptake through the same portal can take place, as shown in Figure 7A and 7C, during the last 8 μs of simulation (light blue color) in S4 (see Table 1). No unique uptake-lipid pathway was detected since the approach to the portals is different for different lipids. The residence time of the lipids inside the cavity is different for every lipid.

**Fig 7.**
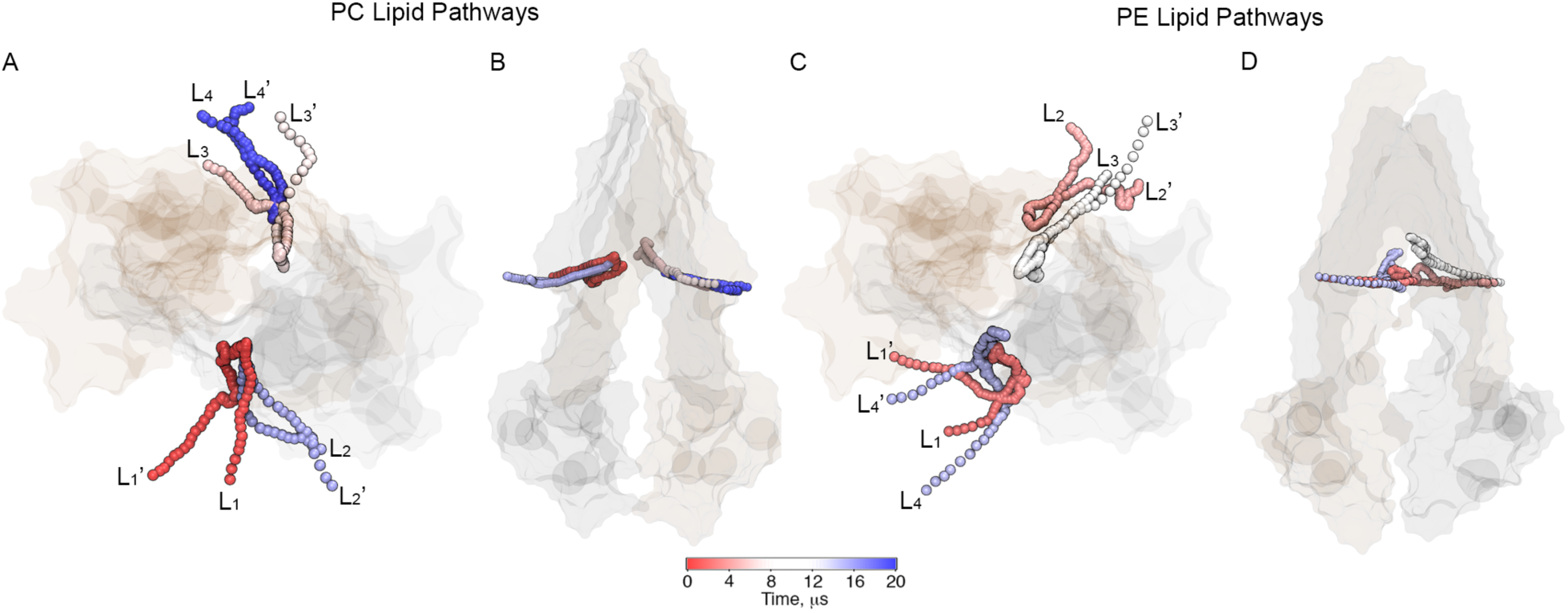
Lipid-uptake pathways found for (A) PC and (B) PE lipid types in plasma membrane ratio PC:PE. Different lipid head groups (PO_4_ beads) access the cavity of P-gp through both portals as a function of time, during 20 μs of simulation time. P-gp is shown in transparent surface representation. Each lipid shown is labeled in the order in which they access the cavity along the simulation by sub index i=[1,…,4]. Before access the cavity, the lipids are labeled as L_i_; after leaving the cavity L_i_’.

## Discussion

P-gp is one of the best characterized and most studied ABC transporters [11][43]. Its broad substrate-specificity allows for the transport of many, chemically diverse molecules [44]–[46], and to date, several studies have suggested poly-specific drug binding sites in the cavity of P-gp [1][45][48][49]. MD simulations have been extensively used to provide molecular details on the interactions between substrates and specific regions of the protein [48], most of them assuming specific binding sites. For example, using umbrella sampling methods, Subramanian *et* al. investigated spontaneous binding for morphine and nicardipine [51]. More recently, Syed *et* al. used the drug binding site in the transmembrane region from docking studies to test interactions with P-gp [52].

Since for P-gp it has been proposed that substrates access the cavity through the membrane [4][51], lipid pathways may also be relevant for elucidating drug transit, uptake and binding. The characterization of lipid binding sites and lipid-uptake mechanisms in P-gp is relevant for a better understanding of drug binding to P-gp and other ABC transporters that are capable of translocating lipids [49][54]. In addition, direct interaction between P-gp and lipid-based molecules as miltefosine and edelfosine, lipid-based anticancer drug molecules, has been previously described [4][42][52]. Towards this aim, we perform CG MD simulations in equilibrium to explore how lipid molecules access the cavity of P-gp. With no previous assumption of specific bindings sites, we identify key residues involved in lipid-uptake, lipid binding and lipid occupancy in the cavity.

## Lipid access to the cavity of P-gp

The structure of P-gp in the inward-facing conformation has two portals (P1, P2) that provide access to the main cavity to molecules located in the lower leaflet of the membrane [22][6]. In all the analyzed simulations, we observed that lipid uptake occurs through these portals for lipids of the inner leaflet, as previously reported by Aller *et* al. [6]. No additional pathways for entrance to the cavity were detected. When comparing the two portals, higher degree of lipid uptake is observed through P1 (Figure 2B). We link this result to the presence of the elbow helix of TMD2, which may screen the interactions between the lipid molecules and the residues located along H12, since it is partially covering H10. This feature also allows the direct exposure of Arg828 of H9 for lipid binding, facing the opening of P2. Although H9 is not included in the standard definition of portals followed in the literature [6][7][22], we found that Arg828 faces P2 and therefore experience lipid binding during the lipid-uptake at P2.

A limitation of the currently available structures of P-gp [22][56][57][58][59] is the missing linker that connects NBD1 with TMD2. This flexible region has been reported as important for the regulation of the substrates specificity and additional effects in the conformational changes in P-gp driven by ATP-ase activity [56][60][61][50]. The presence of this linker may modify the substrate uptake pathways [9], although we believe it is unlikely this linker will affect the lipid binding sites reported in this study.

## Key binding residues for lipid-uptake in P-gp

### At the portals

Our findings suggest that PC and PE lipids access the internal drug-binding pocket through the portals P1 and P2, which has also been reported for drug molecules in several studies [1][4]. It has been also suggested that a positively charged ring of residues such as Arg and Lys [63] [64], could drive substrates towards the cavity, and promote lipid-protein interactions in ABC transporters in general via electrostatic interactions. From our simulations, two binding hot spots at P1 were identified: Lys230 (H4), and Arg355 (H6). This result is in perfect agreement with the experimental report from Marcoux *et* al. [10], where electrostatic interactions between the phosphate groups of lipid molecules and Lys230 where identified by mass spectrometry techniques. Lipid uptake events were also observed at P2, with key binding residues Lys873 (H10) and Arg828 (H9). This set of positively charged residues are key elements for the lipid-uptake due to the high affinity of Lys and Arg for phospholipids head groups via electrostatic interactions, promoting lipid entry into to the cavity.

### In the cavity

The funnel-shaped cavity of P-gp is lined by helices H1/3/5 of TMD1 and H7/11/12 of TMD2. Hydrophobic and aromatic residues in the inner cavity are important for substrate-binding [6][3][5][65][66]. Because of the amphipathic nature of lipids, we investigated the interactions between protein and lipid head group (PO_4_ beads) and between protein and lipid tails. The PO_4_ beads, once inside the cavity, establish interactions mainly with residues of H11 and H12 (Fig 2C and Fig 4B). This result may reflect the structural architecture of the cavity of P-gp. The helical structure of H12 (residues 970-990) is interrupted by a second *α*-helix (residues 991-994), therefore the upper segment of H12 also belongs to the cavity. From our simulations, we observed interactions between the lipid head group and Phe938 on H11 (Figure 4B and SI2), as reported by Aller *et* al. [6] for the inhibitor verapamil. Additional interactions were found also for Ser989, Phe990, Pro992, Val984, Val987 and Ser 988 on H12 and Thr941 Lys929, Lys 930, Met928 and Arg925 on H11 (Figure 4B). In the cavity of P-gp, our findings show the key role of Phe339 as a lipid binding residue. This residue has also been shown to be involved in the binding of ligands such as the cyclic hexapeptide inhibitors, cyclic-tris-(R)-valineselenazole (QZ59-RRR) and cyclic-tris-(S)-valineselenazole (QZ59-SSS) [22], while Zhang *et* al., [67] corroborated the importance of Phe339 for doxorubicin access. If H12 drives many of the interactions between lipid head groups and residues in the cavity, H5 appears to be the key player for the interactions between residues in the cavity of P-gp and the lipid tails (Figure SI3). Residues of H5 such as Leu300, Ile302 and Tyr303 were also reported as binding sites for the cyclic hexapeptide inhibitors, cyclic-tris-(R)-valineselenazole (QZ59-RRR), cyclic-tris-(S)-valineselenazole (QZ59-SSS) and verapamil [6]. Overall, in the case of lipids, the tails extend towards the upper and more hydrophobic region of the cavity, not explored by the PO_4_ beads (Figure 5B,D and Figure 6), with residues such as Phe299, Ile302, Tyr303 and Tyr306 mostly involved in the interactions with the oleoyl tail (Figure SI3). The location of substrates in the top of the cavity has been found in several experimental studies [6][67][68] in agreement with our simulations. Combined, these results reinforce the crucial role of H11, H12 and H5 in the interactions between protein and lipids as substrates.

## Conclusions

In this work we address the lipid-uptake pathways of P-gp in the inward facing conformation in bilayers with different PC:PE lipid ratios. By performing equilibrium MD simulations on a time scale of microseconds, we show that the lipid binding events in the cavity and at the portals of P-gp are accessible and measurable. This approach reveals that uptake events involve lipids from the lower leaflet only, and identifies key lipid binding sites at the portals and inside the cavity of P-gp. In the present study find no selectivity for PC vs. PE lipid-uptake or lipid-binding in P-gp. Further studies focus on substrate release will consider the outward-facing conformation of P-gp for a better understanding of the multidrug efflux pump mechanism.

## Acknowledgements

This work was supported by the Canadian Institutes of Health Research. Additional support came from Alberta Innovates Health Solutions (AIHS) and Alberta Innovates Technology Futures (AITF). R.X.G. is supported by fellowships from AIHS and the Canadian Institutes for Health Research (funding reference number: MFE-140949). DPT is an AIHS Scientist and AITF Strategic Chair in (Bio)Molecular Simulation. Simulations were run on Compute Canada machines, supported by the Canada Foundation for Innovation and partners. This work was undertaken, in part, thanks to funding from the Canada Research Chairs program.

